# Origins and consequences of kinetoplast loss in trypanosomes

**DOI:** 10.64898/2025.12.24.696353

**Authors:** Melanie Ridgway, Douglas O Escrivani, Markéta Novotná, Amy Wood, Michele Tinti, Achim Schnaufer, David Horn

## Abstract

The kinetoplast is the large mitochondrial genome present in the eponymous Kinetoplastida. *Trypanosoma brucei* is an African trypanosome that can lose kinetoplast DNA (kDNA), however, when the nuclear-encoded gamma subunit of the mitochondrial F_1_F_O_-ATP synthase (γATPase) is mutated. These mutations, analogous to a broken camshaft at the core of the ATP synthase rotary motor, are associated with multidrug resistance, and correlated with tsetse-fly independent mechanical transmission, and geographical spread of these parasites beyond Africa. Here we engineer kinetoplast-independent *T. brucei* to explore origins and consequences of kDNA loss. We used oligo targeting to edit the native *γATPase* gene, and selection with the ATP synthase targeting drug oligomycin to enrich the desired mutants. Using this approach, we identified novel M^282^F, M^282^W, and M^282^Y mutants, and subsequently generated precision-edited strains expressing the previously described L^262^P or A^273^P mutants, or the novel M^282^F mutant. Heterozygous M^282^F mutants retained sensitivity to the kDNA-targeting drug acriflavine, while homozygous M^282^F mutants were acriflavine resistant and readily tolerated acriflavine-induced kDNA loss. Proteomics analysis of the homozygous mutant pre-kDNA-loss revealed highly specific depletion of ATP synthase-associated proteins, but not the F_1_ subunits. Complete kDNA-loss in these cells was associated with substantial depletion of kDNA-binding proteins and mitochondrial RNA-processing factors. In contrast, mitochondrial membrane-associated transporters were increased in abundance. We conclude that *T. brucei* cells with a homozygous *γATPase* M^282^F mutation assemble a remodelled ATP synthase and readily tolerate kDNA loss, which is accompanied by substantial remodelling of the mitochondrial proteome

**Author summary:** Mutations in the gamma subunit of the mitochondrial ATP synthase in parasitic African trypanosomes can have major consequences. Specifically, the entire large and complex mitochondrial genome, the kinetoplast, is rendered dispensable, and the cells become resistant to important kinetoplast-targeting drugs. Veterinary parasites with these mutations have also spread outside Africa through simple mechanical transmission, either sexually or by biting flies or vampire bats. We precision-edited the gamma subunit to replicate previously described mutants and identified a novel mutant that readily tolerated kinetoplast loss. Using quantitative proteomics, we demonstrated highly specific depletion of ATP synthase-associated proteins pre-kinetoplast-loss. We then use genome sequencing to show that the kinetoplast could be completely lost by these cells and demonstrated that cells lacking mitochondrial nucleic acids displayed specific depletion of mitochondrial nucleic acid-binding proteins. Notably, several mitochondrial membrane-associated transporter complexes were increased in abundance. Thus, we establish a method to test precise γATPase mutations and to identify new mutations associated with kinetoplast loss. We also show that trypanosomes with a dispensable kinetoplast specifically remodel the ATP synthase pre-kinetoplast-loss and substantially remodel the mitochondrial proteome post-kinetoplast-loss.

## Introduction

*Trypanosoma brucei brucei* is an African trypanosome that is transmitted by tsetse flies, causing nagana disease in cattle and other livestock. Closely related and similarly transmitted African trypanosomes cause sleeping sickness in humans. These parasites are kinetoplastids, flagellated protozoa that contain their mitochondrial genome (mtDNA) in a kinetoplast, hence called kinetoplast DNA, or kDNA. The kDNA is a cytologically prominent feature and comprises a huge network of approx. twenty-five maxicircles and thousands of minicircles, encoding eighteen protein subunits of the mitochondrial respiratory chain, the F_1_F_O_-ATP synthase and the mitoribosome, as well as ribosomal RNA and RNA-editing associated guide RNAs [1, 2]. Insect stage *T. brucei* depend on kDNA-encoded proteins for oxidative phosphorylation, while the bloodstream stage requires F_1_F_O_-ATPase activity to generate the mitochondrial membrane potential; whereby ATP hydrolysis by F_1_ is coupled to proton transfer by F_O_ [3]. Key to this coupling is the central F_1_ γ subunit (γATPase), which acts like a camshaft. γATPase is mechanically coupled to the membrane embedded ring composed of 10 *c* subunits that, together with the A6 subunit (also known as the *a* subunit), forms the proton translocating part of F_O_ [4, 5]. A6 is the only F_1_F_O_-ATPase subunit encoded in kDNA. Because of its essentiality to African trypanosomes and its unique properties, kDNA has proven to be an excellent drug target, albeit with challenges associated with resistance [6, 7].

Remarkably, African trypanosomes of the *T. brucei* group can lose their kDNA, and *T. b. equiperdum and T. b. evansi* present two examples [8, 9]. These parasites still infect equids and various mammals, respectively, and grow as bloodstream forms [7] but cannot differentiate to tsetse insect stages [10]. Although unable to complete the usual life cycle in tsetse, they are transmitted mechanically, either sexually in equids (*equiperdum*), or by biting flies or vampire bats (*evansi*). Consequently, *T. b. equiperdum* and *T. b. evansi* cause diseases known as dourine and surra, respectively, that have spread beyond tsetse endemic regions in Africa, extending to Asia, South America and parts of Europe [11].

Kinetoplast dispensability is caused by specific mutations in γATPase, by allowing for generation of a mitochondrial membrane potential, albeit reduced, in the absence of the kDNA-encoded F_O_ subunit A6 [12–14]. This is thought to involve mechanical uncoupling of the F_1_ and F_O_ components, enhanced ATP hydrolysis by F_1_ and electrogenic exchange of mitochondrial ADP^3-^ for cytosolic ATP^4-^ by the mitochondrial ADP/ATP carrier [15]. Remarkably, beyond the A6 subunit, and the mitoribosome subunits required for its translation, no other kinetoplast-encoded protein appears to be specifically required to maintain the viability of wild-type bloodstream-form *T. b. brucei*. Trypanosomes with γATPase mutations have emerged several times independently, being equivalent to mutations that enable mtDNA loss in petite-negative yeast [8, 14, 15]. These parasites are either dyskinetoplastic or akinetoplastic, lacking some or all of their kDNA, respectively. Other subsequent changes appear to have facilitated adaptation to a tsetse fly independent life-cycle [9].

The kinetoplast has proven to be an excellent drug target, and several veterinary anti-trypanosomal drugs target kDNA, including ethidium bromide and isometamidium. kDNA loss or dispensability in *T. b. evansi, T. b. equiperdum*, and other γATPase mutants renders these cells multi drug resistant, however [7]. Despite connections to the parasite life cycle, geographical disease distribution, and drug resistance, the mechanisms linking γATPase mutations to kDNA dispensability are not fully understood. To develop our understanding of the origins and consequences of kinetoplast loss in trypanosomes, we used oligo-targeting [16] and engineered kinetoplast-independent kinetoplastids. We introduced novel and known mutations in the native *γATPase* gene and found that a novel homozygous M^282^F edit was required to generate kinetoplast-negative strains. We then used proteomics analysis to assess the complement of proteins impacted by γATPase mutation pre- and post-kinetoplast loss, revealing specific impacts on the mitochondrial ATP synthase, mitochondrial nucleic acid binding proteins and mitochondrial membrane-associated transporters.

## Results

### γATPase editing yielded known and novel oligomycin-resistant mutants

Three distinct non-synonymous substitutions have been identified in the γATPase subunit in trypanosomes that display kinetoplast (kDNA) dispensability; L^262^P, A^273^P, and M^282^L [12–14]. A^273^P was identified as a homozygous single-nucleotide mutation in *T. b. equiperdum*, M^282^L was identified as a heterozygous single-nucleotide mutation in some isolates of *T. b. evansi* [14], and L^262^P was later identified as a homozygous single-nucleotide mutation in *T. b. brucei* following acriflavine-selection in the laboratory [12]. The link to kDNA dispensability was validated for both the L^262^P and A^273^P mutations using ectopic *γATPase* expression in transgenic *T. b. brucei*, but a similar assay failed to validate the M^282^L mutation [12].

To assess the impact of specific mutations at the native *γATPase* gene locus (Tb927.10.180) in *T. b. brucei*, we used oligo targeting for precision editing [16], followed by oligomycin selection to enrich those mutants that become independent of the F_O_ component of the ATPase (Fig. 1A); oligomycin targets the proton-binding F_O_ subunit *c* [17]. Given uncertainty regarding the impact of the M^282^L mutation, we began by assessing edits at this site. For oligo targeting, we typically deliver approx. 50 base ‘reverse-strand’ single-stranded oligodeoxynucleotides (ssODNs) by electroporation, and in this case, we designed an ssODN to target the M^282^ site with a centrally located and degenerate ‘NNN’ (N=A, C, T, G) codon (Supplementary Data 1). Wild type *T. b. brucei* cells were transfected in duplicate and grown with oligomycin at 200 nM; approx. three times the EC_50_ (Effective Concentration of drug to inhibit growth by 50 %). We then extracted genomic DNA from surviving cells after six days, PCR-amplified the edited region in the *γATPase* gene, deep-sequenced the *γATPase* amplicons (Fig. 1A), and quantified variant codons. The heatmap in Figure 1B shows relative representation of alternative codons at the targeted site and at flanking sites following oligomycin selection. The analysis revealed highly specific editing at the targeted site, and multiple M^282^ *γATPase* edits enriched in the oligomycin-resistant population, all of which encode aromatic residues, M^282^F, M^282^W, and M^282^Y (Fig. 1B). Notably, all of the enriched mutants required double or triple nucleotide edits while the naturally occurring M^282^L mutation of uncertain significance, accessible via two distinct single nucleotide edits, or four distinct double nucleotide edits, was not enriched.

**Fig. 1:**
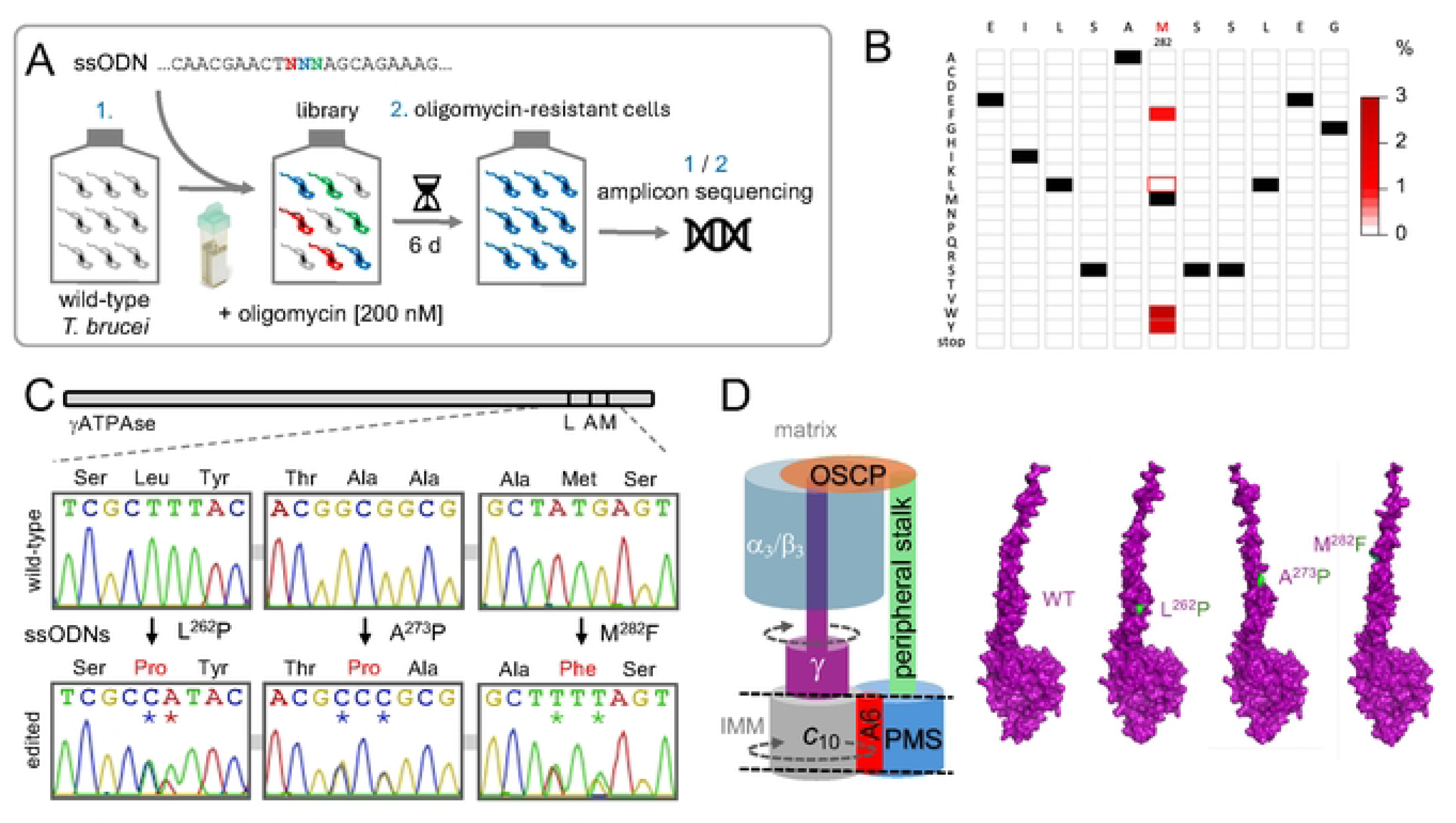
γATPase editing yielded known and novel oligomycin-resistant mutants. **A** The schematic illustrates oligo targeting for saturation mutagenesis of the *T. b*. *brucei γATPase* M^282^ residue. A sixty-four fold degenerate ssODN was transfected into wild-type *T. b. brucei* cells, followed by oligomycin selection and *γATPase* amplicon-sequencing. **B** The heat map shows relative representation of each possible amino acid variant at the targeted M^282^ site and at adjacent sites; averages for two independent oligomycin-resistant cultures relative to an unedited control. More than 8 M reads were mapped per site on average. Unedited codons are indicated (black) as is the previously reported M^282^L mutation. **C** The Sanger sequencing traces show single allele edits encoding the L^262^P, A^273^P and M^282^F mutations, each involving a double nucleotide edit; edited nucleotides are indicated by asterisks. **D** Simplified schematic of the trypanosome F_1_F_O_-ATP synthase and key components discussed here. The α/β hexamer is held in place by attachment to the peripheral ‘stator’ stalk via the OSCP subunit. PMS, peripheral membrane subcomplex; IMM, inner mitochondrial membrane. The AlphaFold models for wild-type (WT) and mutant γATPase were generated using the AlphaFold 3 server, showing mutant residues in green.

We next designed specific ssODNs to introduce one of the novel edits identified above, M^282^F^TTT^, and the other non-synonymous substitutions previously linked to kDNA dispensability, L^262^P^CCA^ and A^273^P^CCC^ (Fig. 1C, Supplementary Data 1); a double nucleotide edit in each case allowed us to distinguish between true edits and spontaneous mutations, the vast majority of which are limited to single nucleotides. We transfected wild type *T. b. brucei* cells with each ssODN, selected the cells with oligomycin at 200 nM, and sub-cloned the resistant cells that emerged. We extracted genomic DNA from the sub-clones, PCR-amplified the edited region in the *γATPase* gene, and Sanger sequenced the amplicons. Sequencing analysis revealed that all three heterozygous edits were effectively introduced (Fig. 1C). We then used AlphaFold3 [18] to visualise how these edits may impact γ-subunit function. The F_1_ component of the ATP synthase comprises a rotor made up of three α-subunits and three β-subunits with a central γ-subunit, which is analogous to a camshaft (Fig. 1D). The models predict conformational defects associated with each γATPase mutation in the extended α-helical camshaft-like segment that may interfere with interaction with the α/β hexamer, perhaps uncoupling F_1_ from F_O_ as described for mitochondrial genome integrity mutations in yeast [19], thereby also reducing sensitivity to the F_O_ inhibitor oligomycin. Thus, *γATPase* precision editing yielded known and novel heterozygous oligomycin-resistant mutants.

### Only bi-allelic M^282^F *γATPase* editing yielded acriflavine-resistant mutants

Prior analyses suggested that γATPase mutant dosage may be important. Specifically, a heterozygous A^281^Δ mutant γATPase allele was reported to be preferentially expressed in *T. b. evansi* [14], while both L^262^P and A^273^P γATPase substitutions with a validated link to kDNA dispensability are present as homozygous mutations in *T. b. brucei* [12] and *T. b. equiperdum* [14], respectively. Since differential expression of mutant alleles could impact the behaviour of heterozygous mutants, we favoured the analysis of homozygous mutants. Although we had not previously observed homozygous editing using oligo targeting, we identified a homozygous *γATPase* M^282^F^TTT^ edited clone following oligomycin selection as detailed above. Indeed, a synonymous polymorphism present seven codons downstream of the targeted codon, allowed us to show that both heterozygous and homozygous M^282^F^TTT^ strains remained diploid at this locus, having retained both *γATPase* alleles (Fig. 2A).

**Fig. 2:**
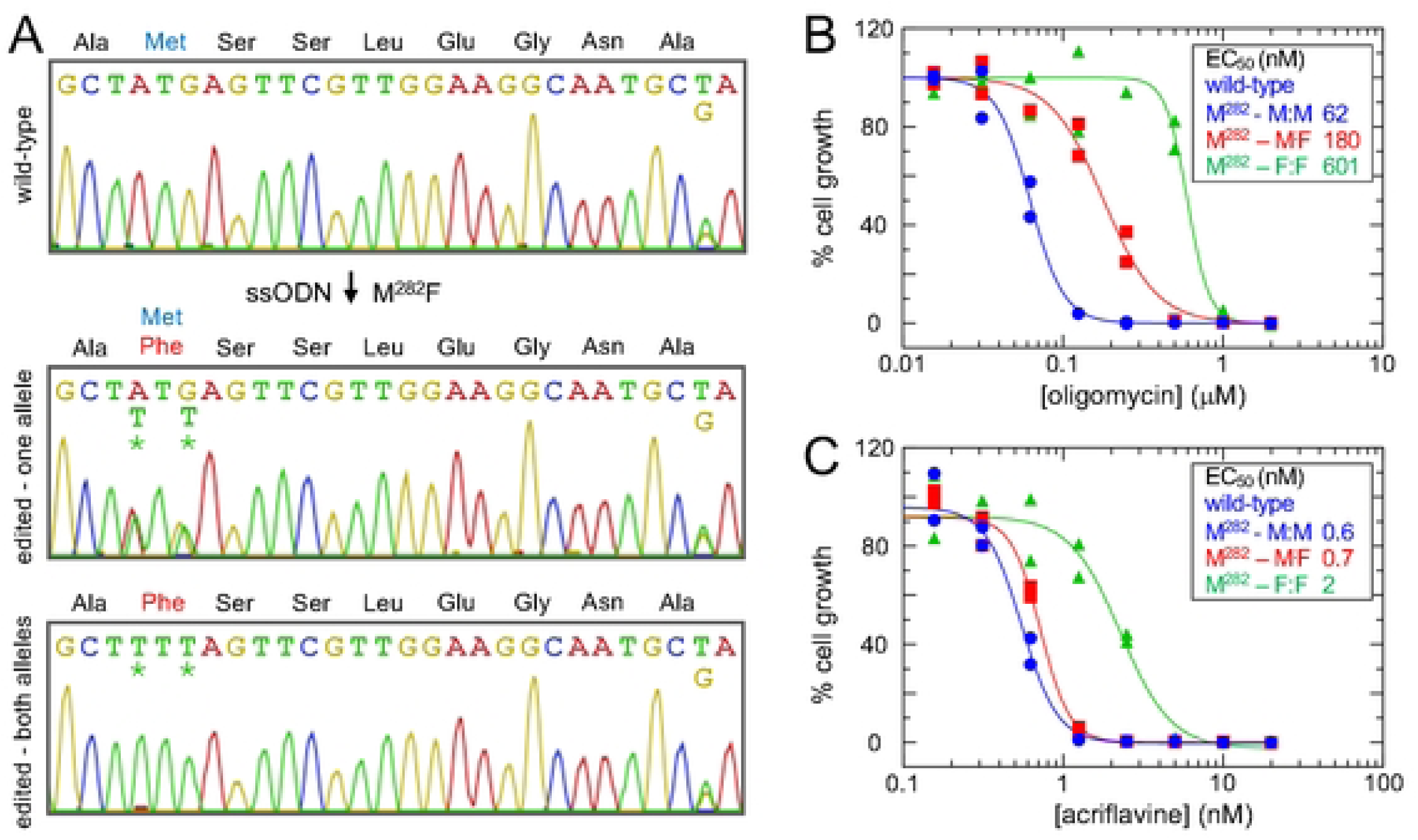
Only bi-allelic *γATPase* editing yielded acriflavine-resistant mutants. **A** The Sanger sequencing traces show *T. b. brucei γATPase* M^282^F edits, both heterozygous and homozygous. A GCT/G, alanine polymorphism can be seen on the right-hand side of each panel, confirming retention of both alleles. Edited nucleotides are indicated by asterisks. **B** Dose-response curves for oligomycin, measured in duplicate. EC_50_ values are shown. **C** Dose-response curves for acriflavine, measured in duplicate. EC_50_ values are shown.

Access to both heterozygous and homozygous M^282^F^TTT^ strains presented an opportunity to compare impacts on oligomycin and acriflavine sensitivity. We performed dose response assays comparing wild-type, heterozygous and homozygous mutants (Fig. 2B-C). An oligomycin dose response assay revealed that heterozygous M^282^F^TTT^ parasites displayed 3-fold increased EC_50_ while homozygous M^282^F^TTT^ mutant parasites displayed 10-fold increased EC_50_ (Fig. 2B); the heterozygous L^262^P and A^273^P mutants displayed 8-fold and 22-fold increased EC_50_, respectively (Supplementary Fig. 1). Thus, all of the *γATPase* edits yielded significant increases in oligomycin resistance (*P* < 1e^−4^), as expected, but the dosage of mutant *γATPase* alleles in the M^282^F^TTT^ edited cells impacted the relative shift in EC_50_.

We next performed dose response assays using acriflavine, which targets kinetoplast DNA, again comparing wild-type, heterozygous and homozygous mutants (Fig. 2C). We observed that homozygous M^282^F^TTT^ parasites displayed a significant, 3-fold increased EC_50_ (*P* < 1e^−4^), while heterozygous M^282^F^TTT^ parasites displayed only 1.1-fold (*P* = 0.4) increase in EC_50_ (Fig. 2C); the heterozygous L^262^P and A^273^P mutants both displayed 2-fold and 2.2-fold increased EC_50_ respectively (Supplementary Fig. 1). Thus, homozygous M^282^F^TTT^ edits yielded acriflavine resistant cells, while heterozygous edits failed to do so, indicating that this mutation is recessive with respect to acriflavine-resistance. We concluded that both heterozygous and homozygous *γATPase* M^282^F^TTT^ editing conferred oligomycin-resistance, albeit to differing degrees, while only homozygous *γATPase* M^282^F^TTT^ editing conferred acriflavine-resistance.

### ATP synthase remodelling and kDNA loss in homozygous *γATPase* mutants

To further elucidate the mechanism underpinning oligomycin and acriflavine cross-resistance, wild-type and homozygous γATPase M^282^F^TTT^ edited parasites were assessed using high resolution quantitative proteomics on an Orbitrap Astral mass spectrometer with data-independent acquisition. The analysis revealed highly specific depletion of all eighteen known nuclear-encoded subunits of the F_O_ component of the *T. b. brucei* ATP synthase (Fig. 3, see Fig. 1D); the peripheral stalk proteins including the oligomycin sensitivity conferring protein OSCP, membrane region proteins and peripheral membrane subcomplex proteins [5]. In striking contrast, subunits of the F_1_ component of the ATP synthase, including the mutated γ subunit itself, were not depleted (Fig. 3). Thus, proteomic analysis revealed highly specific depletion of subunits of the F_O_ component of the *T. b. brucei* ATP synthase in homozygous M^282^F^TTT^ mutants pre kDNA loss.

**Fig. 3:**
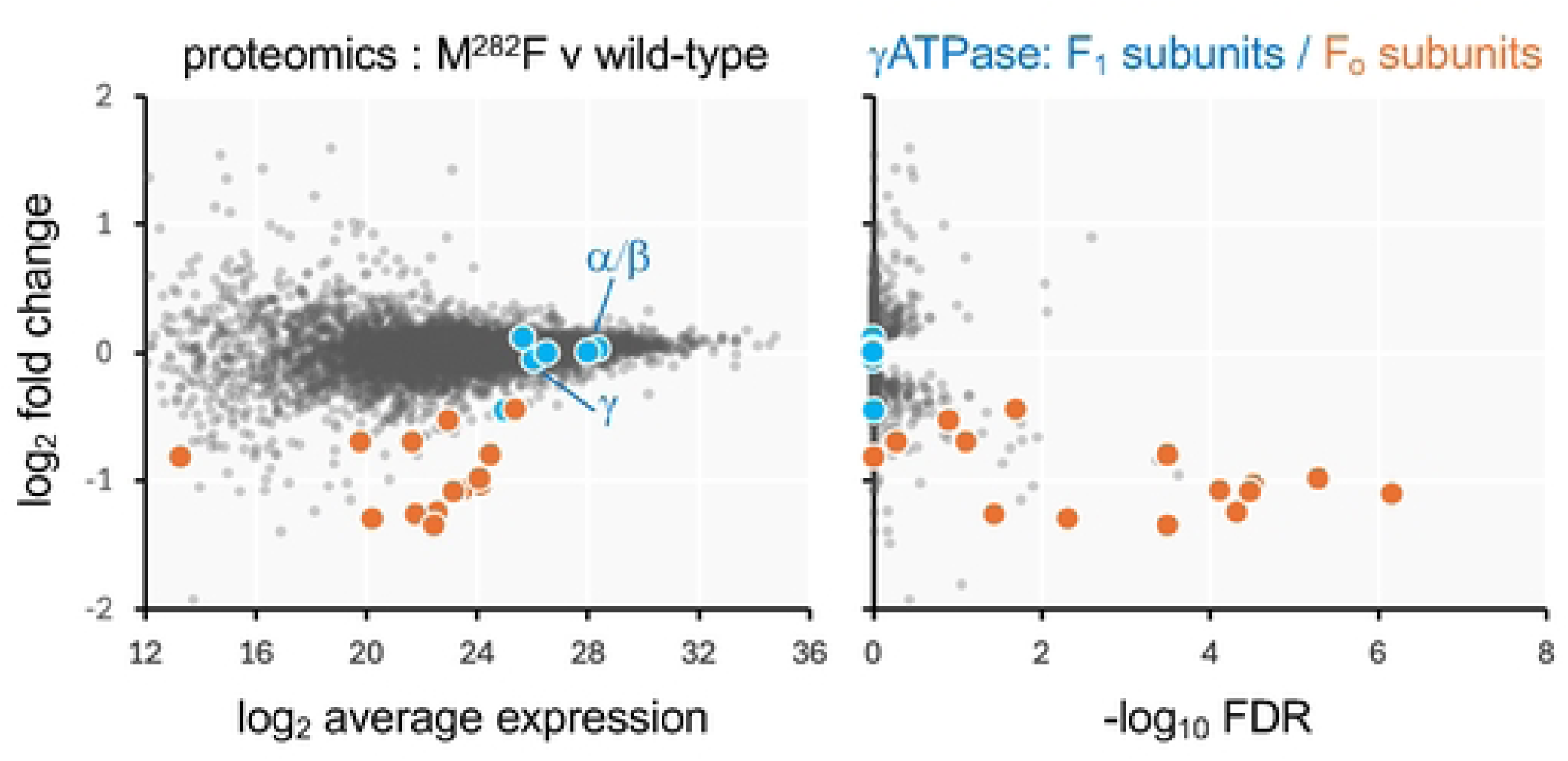
ATP synthase remodelling in homozygous *γATPase* mutants. Proteomics analysis of wild-type *T. b. brucei* and homozygous γATPase M^282^F mutants. Subunits of the F_1_ and F_o_ γATPase components are highlighted. Averages from three replicates; n = 6564.

Acriflavine resistance displayed by homozygous *γATPase* M^282^F^TTT^ edited parasites suggested that kDNA would be dispensable in these cells. To induce kDNA loss, we grew two parallel cultures of homozygous M^282^F^TTT^ edited cells in the presence of a sublethal dose of acriflavine for 7 days (1.25 nM; the EC_50_ is 2 nM, see Fig. 2C). Each population was then cloned by limiting dilution in the absence of acriflavine and clones were assessed by DNA-staining and microscopy. More than 99 % of these cells were kDNA positive prior to exposure to acriflavine and clones that appeared to lack kDNA, two from each independent culture, were selected for further analysis. DNA staining followed by super resolution microscopy of wild-type and kDNA negative cells revealed apparent complete loss of kDNA in the M^282^F^TTT^ mutants (Fig. 4A). Notably, kDNA negative M^282^F^TTT^ cells displayed a growth defect relative to the kDNA positive parent, with doubling time increased by 47 +/-24 % (n = 4); parent doubling time was 6.4 h.

**Fig. 4:**
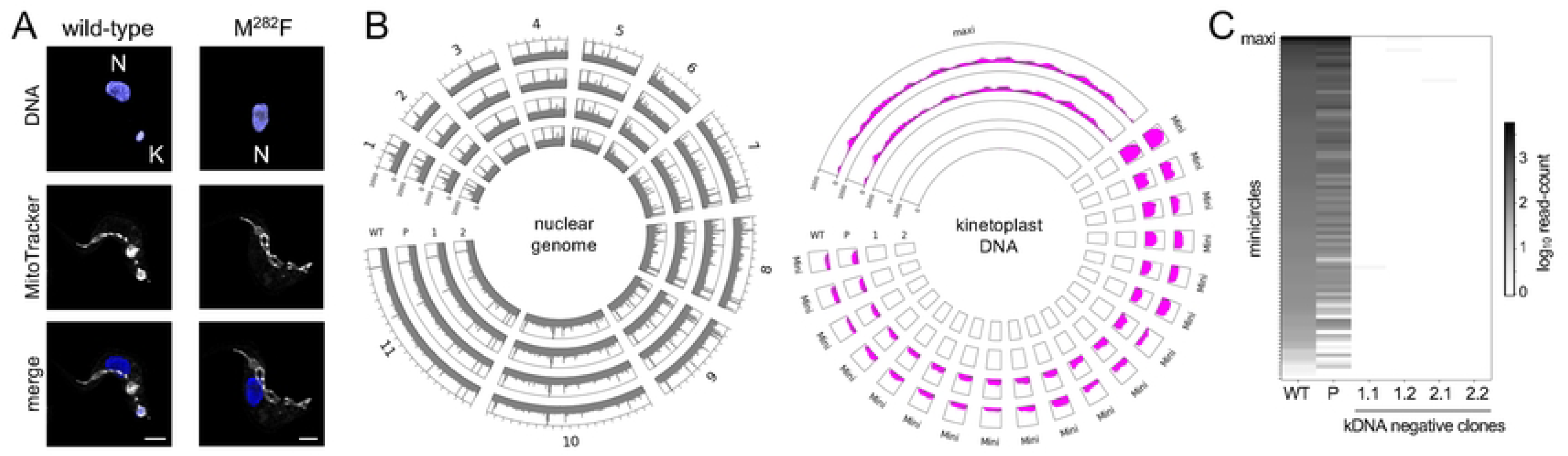
kDNA loss in homozygous *γATPase* mutants. **A** The representative super resolution microscopy images show wild-type *T. b. brucei* and a homozygous γATPase M^282^F mutant lacking kDNA, following growth in acriflavine. DNA was stained with DAPI and mitochondria were stained with MitoTracker. Nuclear DNA (N) and kDNA (K) are indicated. Scale bars, 2 μm. **B-C** Whole genome sequencing data for wild-type (WT) *T. b. brucei*, the homozygous γATPase M^282^F mutant with kDNA (P for parent) and independently generated clones lacking kDNA. **B** The circular plot on the left-hand side shows genome sequencing data mapped to the *T. b. brucei* nuclear chromosomes 1-11 (grey). The circular plot on the right-hand side shows genome sequencing data mapped to the *T. b. brucei* kDNA (magenta), maxicircle sequence and the most abundant minicircle sequences. Mapping is for 150-bp bins and for two independently generated kDNA negative clones. **C** The heatmap shows data for maxicircle sequence, additional minicircle sequences (n = 90), and now for all four kDNA negative clones.

We next considered a more sensitive approach to determine whether kDNA had been completely lost, and subjected wild-type cells, kDNA positive M^282^F^TTT^ edited cells, and all four kDNA negative clones, to whole genome sequencing. We used a recent *T. b. brucei* maxicircle and minicircle DNA assembly for the same strain used in our study [2] as a template for this analysis and observed highly specific loss of both classes of kDNA in independently generated kDNA negative clones without apparent changes in the nuclear genome (Fig. 4B). Closer inspection of minicircle abundance indicated that some were already depleted in M^282^F^TTT^ edited cells prior to acriflavine exposure, with some of the lower abundance minicircles apparently lost entirely (Fig. 4C) Thus, genome sequencing revealed some minicircle loss in homozygous M^282^F^TTT^ mutants pre acriflavine-exposure, and complete elimination of kDNA induced by acriflavine in these cells.

### Mitochondrial proteome remodelling following kDNA loss

To explore the consequences of kDNA loss, we again used high resolution quantitative proteomics to compare homozygous γATPase M^282^F^TTT^ mutants with or without kDNA. We initially examined changes in the abundance of nuclear and mitochondrial proteins following kDNA loss and found that mitochondrial proteins were selectively and significantly reduced in abundance in all four kDNA negative clones described above (Fig. 5A). More detailed analysis of the full proteome revealed substantial changes, including further specific depletion of subunits of the F_O_ component of the *T. b. brucei* ATP synthase, except for subunit *c*, which displayed increased abundance (Fig. 5B). More proteins reported a highly significant (-log_10_ False Discovery Rate [FDR] >4) reduction in abundance relative to proteins that reported increased abundance following kDNA loss (168 v 51), and a closer inspection of >2-fold depleted proteins revealed kDNA-binding proteins and mitochondrial RNA-processing factors (Fig. 5C), consistent with destabilisation of these proteins after loss of all mitochondrial DNA and RNA. These included the kDNA-associated proteins involved in DNA compaction [20], primases [21] polymerases [22], topoisomerase involved in DNA replication, and mRNA polyadenylation factors [23]. Other notable depleted proteins were the calcium uniporter, known to interact with subunit *c* of the ATP-synthase [24], and PUF9, an RNA-binding protein involved in cell cycle regulation [25]. Notably, IF1, an inhibitor of F_1_-mediated ATP hydrolysis, was also depleted. Expression of this protein was thought to be restricted to the insect stage, where ATP synthesis, but not hydrolysis, is essential [26]. This observation might indicate increased ATP hydrolysis following kDNA loss. In contrast, PUF9 target 1 (PNT1), a kDNA replication-associated peptidase [27], is notably increased in abundance in kDNA negative cells (Fig. 5C).

**Fig. 5:**
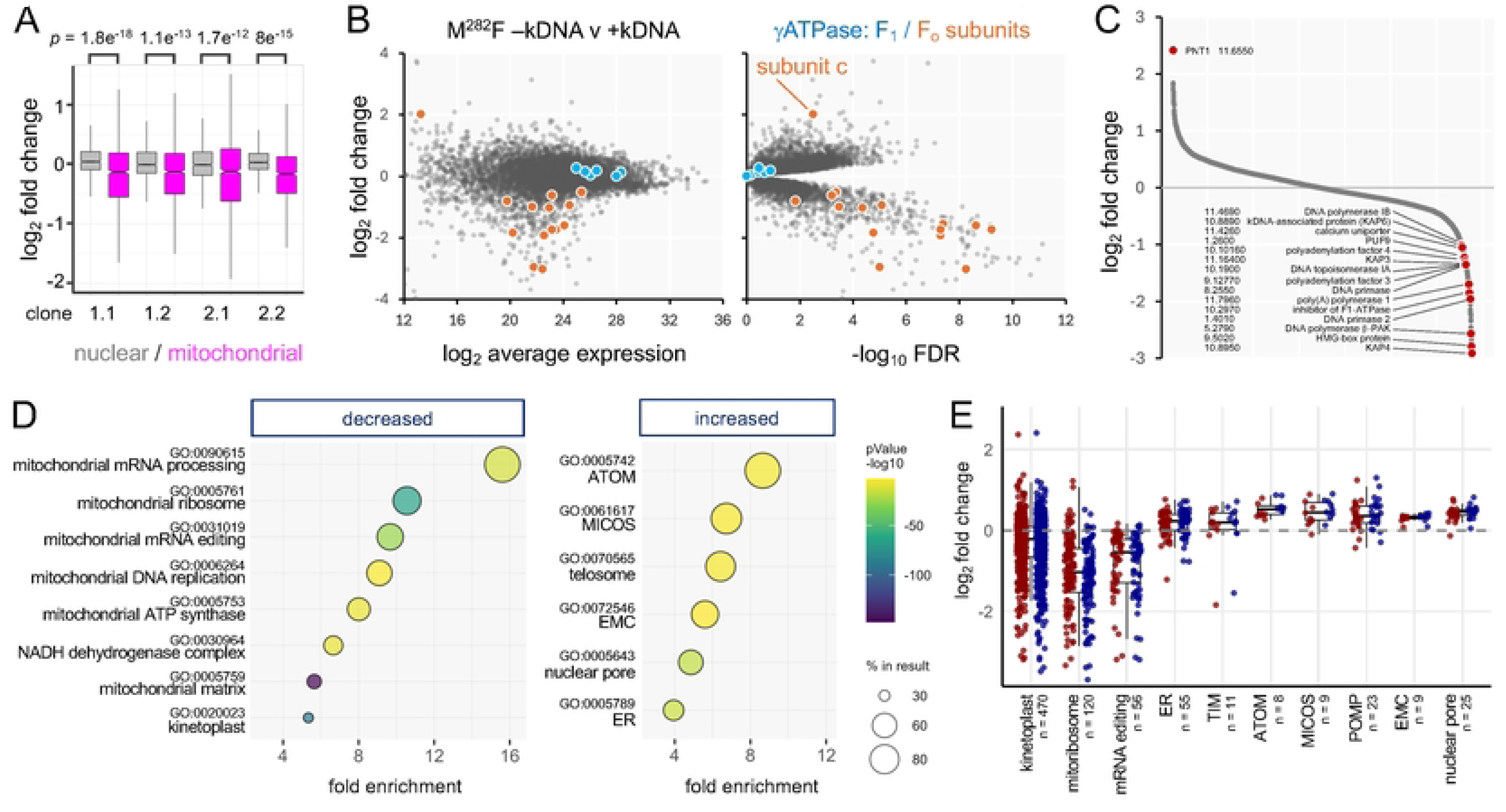
Mitochondrial proteome remodelling following kDNA loss. **A** Proteomics analysis of homozygous γATPase M^282^F *T. b. brucei* mutants and all four M^282^F clones lacking kDNA. The boxplot shows nuclear proteins (n = 1185) and mitochondrial proteins (n = 1431). Boxes indicate the interquartile range (IQR) and the whiskers show the range of values within 1.5 × IQR. **B** Proteomics analysis for clone 1.2 with subunits of the F_1_ and F_o_ γATPase components highlighted. Averages from three replicates; n = 6768. **C** Proteomics analysis as in B showing all proteins with log_2_ average expression >16 and with some notable proteins highlighted. **D** Gene Ontology profiles for proteins with log_2_ average expression >16 that are significantly (>2 –log_10_ FDR) decreased or increased in abundance in kDNA negative cells. **E** Selected cohorts of proteins that are significantly decreased or increased in abundance in kDNA negative cells. Boxes indicate the interquartile range (IQR) and the whiskers show the range of values within 1.5 × IQR. ER, Endoplasmic Reticulum; TIM, Translocases of the Inner Membrane; ATOM, Archaic Translocase of the Outer Membrane; MICOS, Mitochondrial contact site and Cristae Organizing System; POMP, Present in the Outer mitochondrial Membrane Proteome; EMC, ER-Membrane Complex. All cohorts derived using GO-terms except for TIM and POMP, derived using wild-card searches, TIM* and POMP* at https://tritrypdb.org.

We next profiled proteins that were significantly (FDR <0.01) reduced (n = 507) or increased (n = 761) in abundance following kDNA loss using Gene Ontology (GO) terms. The top GO-term hits for depleted proteins were exclusively associated with the mitochondrion, again including DNA and RNA binding proteins, and ATP synthase (Fig. 5D). Kinetoplast (*P* = 1.4e^−89^) and mitochondrial matrix (*P* = 4.1e^−148^) registered the highest significance, and components of the NADH dehydrogenase, complex I of the electron transport chain, were also significantly depleted; perhaps unsurprisingly since several components of this complex are encoded in kDNA [1, 2]. In contrast, mitochondrial membrane-associated transporters, and endoplasmic reticulum (ER) associated proteins, were significantly increased in abundance. The telomere-telomerase complex and nuclear pore proteins were also increased, perhaps reflecting connections between nuclear and kDNA replication [28]. Changes in abundance for several of these cohorts of proteins are shown in Figure 5E, revealing substantial depletion of the mitochondrial ribosome and RNA editing complex [29]. The archaic translocase of the outer mitochondrial membrane (ATOM), translocase of the inner mitochondrial membrane (TIM) [30], and mitochondrial contact site and cristae organization system [31], were all increased in abundance (Fig. 5E), perhaps compensating for mitochondrial import defects associated with reduced mitochondrial membrane potential [13, 32]. Proteins present in the outer mitochondrial membrane proteome (POMP), many of which remain otherwise uncharacterised [33], were also increased in abundance. Finally, the ER-membrane complex (EMC), previously connected to kinetoplast dependency [6], and now known to localise to the mitochondrial – ER interface [34], was also increased in abundance (Fig. 5E). Taken together, our results reveal ATP synthase complex remodelling associated with bi-allelic *γATPase* mutation and multi-drug resistance. These cells readily tolerate kDNA loss, which is associated with substantial mitochondrial proteome remodelling.

## Discussion

γATPase mutations in *T. brucei* are associated with kinetoplast DNA loss and multidrug resistance and are also correlated with tsetse-fly independent mechanical transmission and geographical spread of these parasites beyond Africa. Here, we explore γATPase mutations and connections to kinetoplast DNA loss in *T. b. brucei*. We precision-edited the native *γATPase* gene, confirming that L^262^P and A^273^P mutants are resistant to the ATP synthase targeting drug oligomycin and to the kDNA-targeting drug, acriflavine. We also identified novel oligomycin-resistant aromatic amino-acid mutants replacing M^282^. Quantitative proteomics analysis of homozygous M^282^ mutants revealed specific depletion of ATP synthase F_O_ components prior to kDNA loss. Following acriflavine-induced kDNA loss, confirmed to be complete by genome sequencing, we observed substantial mitochondrial proteome remodelling; the abundance of mitochondrial DNA and mRNA binding proteins was reduced while the abundance of proteins involved in mitochondrial import was increased. While we observed further depletion of ATP synthase F_O_ components following kDNA loss, *c* subunit abundance was increased, perhaps reflecting accumulation of F_1_-ATP synthase associated with only the *c*-ring of the F_O_ moiety (see Fig. 1D).

In terms of the origins of kDNA loss, we found that precision-editing could be used to generate both heterozygous and homozygous γATPase mutants in otherwise wild-type trypanosomes. While a heterozygous M^282^F edit failed to confer resistance to acriflavine, a homozygous M^282^F mutant was acriflavine-resistant and readily tolerated kDNA loss. We used quantitative proteomics to explore the impact of the homozygous M^282^F edit prior to kDNA loss and observed highly specific depletion of ATP synthase-associated proteins but not the F_1_ subunits. Importantly, we selected for edited cells using oligomycin rather than a DNA-damaging agent, avoiding direct selective pressure on the kDNA and likely reducing the potential for off-target mutations at this stage of the process. It is also worth noting in this regard that the widespread use of veterinary anti-trypanosomal drugs that target the kDNA, such as the DNA-damaging agents, ethidium bromide and isometamidium, could induce γATPase mutations and/or other mutations [35].

A homozygous M^282^F γATPase edit generated here in *T. b. brucei* was sufficient to confer kDNA dispensability, while a heterozygous edit was insufficient. This recessive effect with respect to kDNA dispensability suggested a dosage effect and prompted further consideration of naturally occurring non-synonymous γATPase mutations implicated in conferring kDNA dispensability; a heterozygous M^282^L mutation in some isolates of *T. b. evansi* [14], a homozygous A^273^P mutation in *T. b. equiperdum* [14], and a homozygous L^262^P mutation in *T. b. brucei* [12]. Among these non-synonymous edits, we only failed to recover M^282^L using oligomycin selection. Indeed, ectopic mutant γATPase expression assays in *T. b. brucei* also indicated that the M^282^L mutation is insufficient to confer kDNA dispensability [12]. On the other hand, similar assays suggested that a single A^273^P or L^262^P allele may be sufficient to confer kDNA dispensability, and our demonstration that heterozygous A^273^P or L^262^P edits are sufficient to confer acriflavine resistance is consistent with this view.Homozygous editing described here is the first example of dual-allele oligo targeting, suggesting that this approach may be exploited to generate and assess further homozygous *γATPase* mutants.

We also used quantitative proteomics to elucidate the consequences of kDNA loss in homozygous M^282^F γATPase mutants and observed extensive proteome remodelling in this case. Perhaps unsurprisingly, kDNA-binding proteins and mitochondrial RNA-processing factors were significantly depleted in the absence of mitochondrial DNA and RNA; the residual editing complexes in kDNA-negative cells have been reported to retain function, however [36]. Mitochondrial membrane-associated transporters on the other hand were significantly increased in abundance, suggesting a boost in mitochondrial protein import capacity and inter-organellar trafficking capacity. These adaptations may partially compensate for mitochondrial import defects associated with reduced mitochondrial membrane potential or may reflect a response to depletion of multiprotein complexes that contain kDNA-encoded proteins, such as respiratory complex I, the F_1_F_O_-ATP synthase and the mitoribosome [13, 32]. The adaptations were not sufficient to recover a wild-type growth rate, however, consistent with further adaptations reported in *T. b. evansi* and*T. b. equiperdum* [9]. One adaptation we found particularly intriguing was increased abundance of components of the ER membrane complex, since we previously linked expression of this complex to kDNA dispensability in the absence of γATPase mutation [6]; this complex is now known to localise to the mitochondrial – ER interface [34]. Also increased in abundance was subunit *c* of the ATP synthase, which interacts with the calcium uniporter [24], although this latter complex was reduced in abundance. Notably, association of mutant F_1_ with the inner mitochondrial membrane, perhaps proximal to the ATP/ADP carrier, is thought to be required to sustain mitochondrial membrane potential following kDNA loss [15, 37]. We suggest that, in the absence of the kDNA-encoded subunit A6, ATP synthase assembly is arrested after attachment of the F_1_ moiety to the *c*-ring, consistent with the assembly pathway elucidated in other eukaryotes [38]. This model is consistent with our observations and is also consistent with detection of putative ‘F1-*c*’ complexes by native gel electrophoresis in dyskinetoplastic *T. brucei* [37]. Other adaptations may reflect loss of prior communication between the kDNA and nuclear DNA. For example, PUF9 target 1, a kDNA replication-associated peptidase [27], was increased in abundance while the RNA-binding protein PUF9, involved in cell cycle regulation [25], was reduced in abundance. The telomere-telomerase complex was also increased in abundance. Taken together, these adaptations reveal a remarkable connectivity between the ATP synthase and other mitochondrial and even other cellular complexes and compartments.

In conclusion, *T. brucei* cells with a bi-allelic *γATPase* defect assemble a remodelled ATP synthase complex, and tolerate kDNA-loss, accompanied by substantial mitochondrial proteome remodelling. Proline mutations with the potential to disrupt helical structure at L^262^ or A^273^ in the γATPase [12, 14], or bulky aromatic residue mutations at M^282^, introduce defects analogous to a broken camshaft at the core of this ATP synthase rotary motor. Our findings yield new insights into the origins and consequences of kinetoplast loss, with implications for the evolution of trypanosome sub-species that have global veterinary impacts.

## Methods

### *T. brucei* growth and *γATPase* gene editing

Bloodstream form *T. b. brucei* Lister 427 cells were grown in HMI-11 (Gibco) supplemented with 10 % fetal bovine serum (Sigma) at 37°C and with 5 % CO_2_ in a humidified incubator. For site saturation mutagenesis using oligo-targeting, a degenerate ssODN was transfected in duplicate by electroporation with a Nucleofector (Lonza), and a human T-cell kit (Lonza), with the Nucleofector set to Z-001 (Amaxa). Briefly, we used 40 µg of the ssODNs in 10 µl of 10 mM Tris-Cl, pH 8.5, mixed with 25 million cells in 100 µl transfection buffer. 200 nM oligomycin was applied 6 h after transfection. DNA was isolated 7 d later. The *γATPase* fragment was amplified by PCR using Q5 high fidelity DNA polymerase (New England Biolabs) as per the manufacturer’s instructions, and primers 1 and 2. Annealing was at 63°C and elongation was for 30 s. PCR products were purified using a Qiagen PCR purification kit. For specific mutagenesis using oligo targeting, a specific ssODN was transfected in duplicate with 10 million cells, and 200 nM oligomycin was applied 6 h after transfection. Oligomycin-resistant cultures were sub-cloned by serial dilution in 96-well plates 5 d later and DNA was extracted from the clones. The *γATPase* fragment was amplified by PCR as above but using primers 1 and 3 in this case. The products were Sanger sequenced using primer 4 at Azenta Life Sciences. To induce kinetoplast loss, cells were grown in the presence of 1.25 nM acriflavine for 7 days and then subcloned.

### Dose-response assays

To determine the Effective Concentration of drug to inhibit growth by 50 % (EC_50_), cells were plated in 96-well plates at 1 × 10^3^ cells/ml in a 2-fold serial dilution of selective drug. Plates were incubated at 37°C for 72 h. 20 µl resazurin sodium salt (AlamarBlue, Sigma) at 0.49 mM in PBS was then added to each well, and plates were incubated for a further 6 h. Fluorescence was determined using an Infinite 200 pro plate reader (Tecan) at an excitation wavelength of 540 nm and an emission wavelength of 590 nm. EC_50_ values were derived using Prism (GraphPad).

### Proteomics

Mass spectrometry and proteomic analyses were performed as described previously [39]. Briefly, triplicate samples of 5 × 10^7^ cells were washed in PBS and resuspended in 100 μL of 5% sodium dodecyl sulphate, 100 mM triethyl ammonium bicarbonate. Samples were analysed on an Astral mass spectrometer at the Fingerprints Proteomics Facility, University of Dundee, by directDIA using Spectronaut software (Biognosys). Data analysis was performed using custom Python and R scripts, using the SciPy ecosystem of open-source software libraries [40]. A protein group pivot table was exported from the output of the Spectronaut analysis v19 (Biognosys). The differential expression analysis was performed with limma v3.54 [41] after log2 transformation of the data. FDR values were computed with the toptable function in limma.

### Genome Sequencing

For analysis following site saturation mutagenesis, PCR amplicons were sequenced at the Beijing Genome Institute (BGI) on a DNBseq platform with 150 base paired-end reads as described previously [16] and codon-based read counts were derived using the OligoSeeker software [42]. Whole genome sequencing data were analysed with alignment to the *T. b. brucei* reference genome v46 clone 427_2018 supplemented with 427 maxi and minicircle sequences [2]. The alignment and read counts were performed with the automated snakemake [43] pipeline myRNA-SEQ [44]. Read coverage was extracted from the BAM files using a bin size of 150 with the bamCoverage function from deepTools (v 3.5.6). The circular visualization was performed with the pyCirclize (1.6) Python package (https://github.com/moshi4/pyCirclize).

### Microscopy

To identify clones lacking kDNA, cells were fixed in 1 % paraformaldehyde (PFA) for 15 min, washed twice in PBS and resuspended in water with 1 % bovine serum albumin (BSA). Cells were attached to a 12-well 5 mm slide (Thermo Scientific) by drying overnight. After rehydration in PBS for 5 min, slides were mounted in Vectashield with DAPI and sealed under a coverslip. Cells were viewed at 63x magnification with oil immersion on a Zeiss Axiovert 200 M microscope with Zen Pro software (Zeiss). For super resolution microscopy, cells were stained with 100 nM Mitotracker red CMXRos (Invitrogen) for 5 min at 37°C prior to fixation in 3 % PFA for 15 min, washed in PBS then resuspended in water with 1 % BSA. Cells were attached to poly-lysine-coated coverslips for 4 h at room temperature then stained with DAPI for 30 min before mounting in Vectashield (without DAPI) and sealing to a glass slide. Cells were imaged as z-stacks (0.1–0.2 μm) at 63x magnification on a Leica Stellaris 8 inverted confocal microscope equipped with Power HyD detectors and subjected to adaptive deconvolution using the integrated Leica LIGHTNING algorithm for super-resolution microscopy. Images were analysed using Fiji v1.5.2e.

## Data availability

High-throughput sequencing data have been deposited in the European Nucleotide Archive, www.ebi.ac.uk/ena (BioProject ID: PRJNA1380964). The mass spectrometry proteomics data have been deposited at the PRIDE repository (Dataset identifier PXD071938).

## Acknowledgements

We thank Gustavo Bravo Ruiz for assistance with visualising GO-term profiles.

## Author Contributions

The oligo targeting experiments were designed by M.R. and D.H. and carried out by M.R. and A.W. Dose response assays were performed by M.R., D.O.E. and M.N. Proteomics and amplicon and genome sequence data analyses were performed by M.T., A.S. and D.H. Super-resolution microscopy was performed by D.O.E. The work was supervised by D.H. The manuscript was written by M.R., D.O.E. and D.H. The manuscript was edited by all authors.

## Competing interests

The authors declare no competing interests.

Supplementary Information is available for this paper.

Correspondence and requests for materials should be addressed to

---

**Supplementary Fig. 1:**
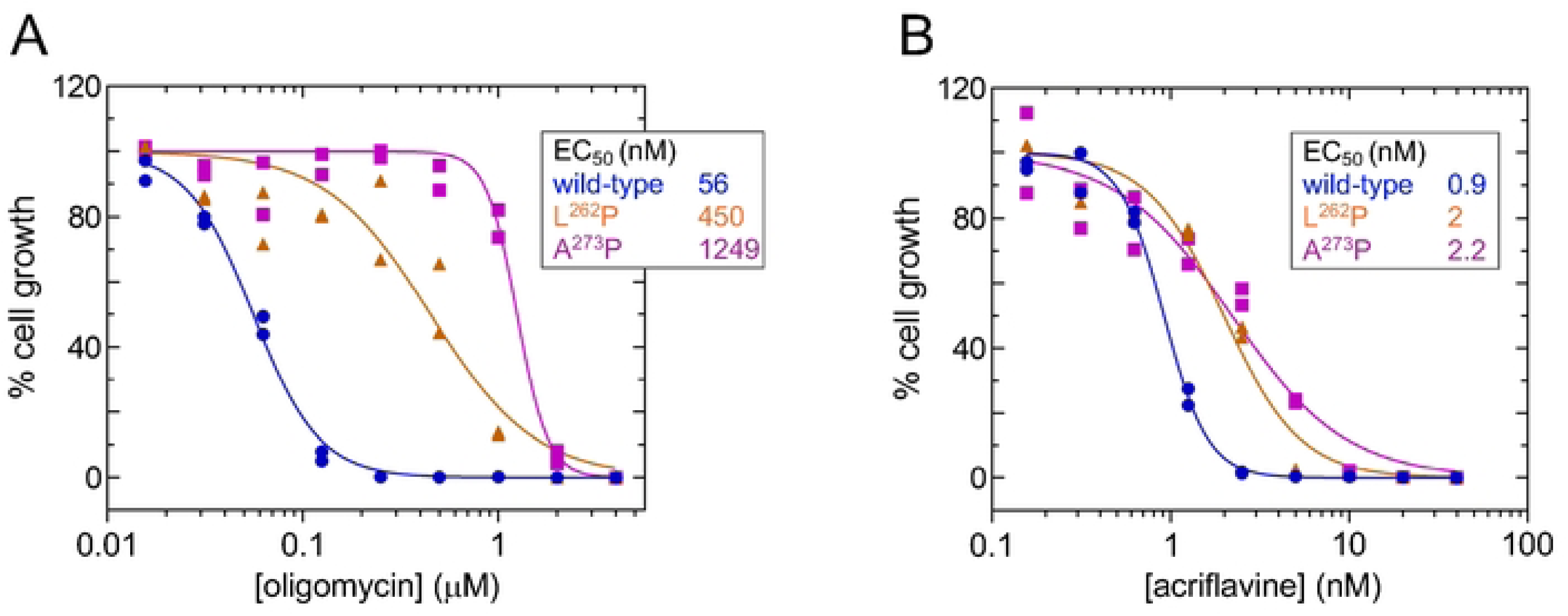
**A** Dose-response curves for oligomycin, measured in duplicated. **B** Dose-response curves for acriflavine, measured in duplicated.

## References

1. Cooper S, Wadsworth ES, Schnaufer A, Savill NJ. Organization of minicircle cassettes and guide RNA genes in *Trypanosoma brucei*. RNA. 2022;28(7):972–92. Epub 2022/04/14. doi: 10.1261/rna.079022.121. PubMed PMID: 35414587; PubMed Central PMCID: PMCPMC9202587.

2. Zhao X, He Y, Zhang F, Aphasizheva I, Aphasizhev R, Zhang L. Comparative mitochondrial genome and transcriptome analyses reveal strain-specific features of RNA editing in *Trypanosoma brucei*. Nucleic Acids Res. 2025;53(13). Epub 2025/07/17. doi: 10.1093/nar/gkaf661. PubMed PMID: 40671526; PubMed Central PMCID: PMCPMC12266143.

3. Zikova A. Mitochondrial adaptations throughout the *Trypanosoma brucei* life cycle. J Eukaryot Microbiol. 2022;69(6):e12911. Epub 2022/03/25. doi: 10.1111/jeu.12911. PubMed PMID: 35325490.

4. Gahura O, Hierro-Yap C, Zikova A. Redesigned and reversed: architectural and functional oddities of the trypanosomal ATP synthase. Parasitology. 2021;148(10):1151–60. Epub 2021/02/09. doi: 10.1017/S0031182021000202. PubMed PMID: 33551002; PubMed Central PMCID: PMCPMC8311965.

5. Gahura O, Muhleip A, Hierro-Yap C, Panicucci B, Jain M, Hollaus D, et al. An ancestral interaction module promotes oligomerization in divergent mitochondrial ATP synthases. Nat Commun. 2022;13(1):5989. Epub 2022/10/12. doi: 10.1038/s41467-022-33588-z. PubMed PMID: 36220811; PubMed Central PMCID: PMCPMC9553925.

6. Baker N, Hamilton G, Wilkes JM, Hutchinson S, Barrett MP, Horn D. Vacuolar ATPase depletion affects mitochondrial ATPase function, kinetoplast dependency, and drug sensitivity in trypanosomes. Proc Natl Acad Sci U S A. 2015;112(29):9112–7 Epub 2015/07/08. doi: 10.1073/pnas.1505411112. PubMed PMID: 26150481; PubMed Central PMCID: PMCPMC4517229.

7. Gould MK, Schnaufer A. Independence from Kinetoplast DNA maintenance and expression is associated with multidrug resistance in *Trypanosoma brucei in vitro*. Antimicrob Agents Chemother. 2014;58(5):2925–8. Epub 2014/02/20. doi: 10.1128/AAC.00122-14. PubMed PMID: 24550326; PubMed Central PMCID: PMCPMC3993240.

8. Carnes J, Anupama A, Balmer O, Jackson A, Lewis M, Brown R, et al. Genome and phylogenetic analyses of *Trypanosoma evansi* reveal extensive similarity to *T. brucei* and multiple independent origins for dyskinetoplasty. PLoS Negl Trop Dis. 2015;9(1):e3404. Epub 2015/01/09. doi: 10.1371/journal.pntd.0003404. PubMed PMID: 25568942; PubMed Central PMCID: PMCPMC4288722.

9. Oldrieve GR, Venter F, Cayla M, Verney M, Hebert L, Geerts M, et al. Mechanisms of life cycle simplification in African trypanosomes. Nat Commun. 2024;15(1):10485. Epub 2024/12/03. doi: 10.1038/s41467-024-54555-w. PubMed PMID: 39622840; PubMed Central PMCID: PMCPMC11612274.

10. Dewar CE, MacGregor P, Cooper S, Gould MK, Matthews KR, Savill NJ, et al. Mitochondrial DNA is critical for longevity and metabolism of transmission stage *Trypanosoma brucei*. PLoS Pathog. 2018;14(7):e1007195. Epub 2018/07/19. doi: 10.1371/journal.ppat.1007195. PubMed PMID: 30020996; PubMed Central PMCID: PMCPMC6066258.

11. Schnaufer A, Domingo GJ, Stuart K. Natural and induced dyskinetoplastic trypanosomatids: how to live without mitochondrial DNA. Int J Parasitol. 2002;32(9):1071–84. Epub 2002/07/16. doi: 10.1016/s0020-7519(02)00020-6. PubMed PMID: 12117490.

12. Dean S, Gould MK, Dewar CE, Schnaufer AC. Single point mutations in ATP synthase compensate for mitochondrial genome loss in trypanosomes. Proc Natl Acad Sci U S A. 2013;110(36):14741–6. Epub 2013/08/21. doi: 10.1073/pnas.1305404110. PubMed PMID: 23959897; PubMed Central PMCID: PMCPMC3767566.

13. Eze AA, Gould MK, Munday JC, Tagoe DN, Stelmanis V, Schnaufer A, et al. Reduced mitochondrial membrane potential is a late adaptation of *Trypanosoma brucei brucei* to isometamidium preceded by mutations in the gamma subunit of the F1Fo-ATPase. PLoS Negl Trop Dis. 2016;10(8):e0004791. Epub 2016/08/16. doi: 10.1371/journal.pntd.0004791. PubMed PMID: 27518185; PubMed Central PMCID: PMCPMC4982688.

14. Lai DH, Hashimi H, Lun ZR, Ayala FJ, Lukes J. Adaptations of *Trypanosoma brucei* to gradual loss of kinetoplast DNA: *Trypanosoma equiperdum* and *Trypanosoma evansi* are petite mutants of *T. brucei*. Proc Natl Acad Sci U S A. 2008;105(6):1999–2004. Epub 2008/02/05. doi: 10.1073/pnas.0711799105. PubMed PMID: 18245376; PubMed Central PMCID: PMCPMC2538871.

15. Schnaufer A, Clark-Walker GD, Steinberg AG, Stuart K. The F1-ATP synthase complex in bloodstream stage trypanosomes has an unusual and essential function. EMBO J. 2005;24(23):4029–40. Epub 2005/11/05. doi: 10.1038/sj.emboj.7600862. PubMed PMID: 16270030; PubMed Central PMCID: PMCPMC1356303.

16. Altmann S, Rico E, Carvalho S, Ridgway M, Trenaman A, Donnelly H, et al. Oligo targeting for profiling drug resistance mutations in the parasitic trypanosomatids. Nucleic Acids Res. 2022;50(14):e79. Epub 2022/05/08. doi: 10.1093/nar/gkac319. PubMed PMID: 35524555; PubMed Central PMCID: PMCPMC9371896.

17. Symersky J, Osowski D, Walters DE, Mueller DM. Oligomycin frames a common drug-binding site in the ATP synthase. Proc Natl Acad Sci U S A. 2012;109(35):13961–5. Epub 2012/08/08. doi: 10.1073/pnas.1207912109. PubMed PMID: 22869738; PubMed Central PMCID: PMCPMC3435195.

18. Jumper J, Evans R, Pritzel A, Green T, Figurnov M, Ronneberger O, et al. Highly accurate protein structure prediction with AlphaFold. Nature. 2021;596(7873):583–9. Epub 2021/07/16. doi: 10.1038/s41586-021-03819-2. PubMed PMID: 34265844; PubMed Central PMCID: PMCPMC8371605.

19. Arsenieva D, Symersky J, Wang Y, Pagadala V, Mueller DM. Crystal structures of mutant forms of the yeast F1 ATPase reveal two modes of uncoupling. J Biol Chem. 2010;285(47):36561–9. Epub 2010/09/17. doi: 10.1074/jbc.M110.174383. PubMed PMID: 20843806; PubMed Central PMCID: PMCPMC2978584.

20. Wang J, Pappas-Brown V, Englund PT, Jensen RE. TbKAP6, a mitochondrial HMG box-containing protein in *Trypanosoma brucei*, is the first trypanosomatid kinetoplast-associated protein essential for kinetoplast DNA replication and maintenance. Eukaryot Cell. 2014;13(7):919–32. Epub 2014/06/01. doi: 10.1128/EC.00260-13. PubMed PMID: 24879122; PubMed Central PMCID: PMCPMC4135736.

21. Hines JC, Ray DS. A second mitochondrial DNA primase is essential for cell growth and kinetoplast minicircle DNA replication in *Trypanosoma brucei*. Eukaryot Cell. 2011;10(3):445–54. Epub 2011/01/25. doi: 10.1128/EC.00308-10. PubMed PMID: 21257796; PubMed Central PMCID: PMCPMC3067476.

22. Delzell SB, Nelson SW, Frost MP, Klingbeil MM. *Trypanosoma brucei* mitochondrial DNA polymerase POLIB contains a novel polymerase domain insertion that confers dominant exonuclease activity. Biochemistry. 2022;61(23):2751–65. Epub 2022/11/19. doi: 10.1021/acs.biochem.2c00392. PubMed PMID: 36399653; PubMed Central PMCID: PMCPMC9731263.

23. Aphasizheva I, Yu T, Suematsu T, Liu Q, Mesitov MV, Yu C, et al. Poly(A) binding KPAF4/5 complex stabilizes kinetoplast mRNAs in *Trypanosoma brucei*. Nucleic Acids Res. 2020;48(15):8645–62. Epub 2020/07/03. doi: 10.1093/nar/gkaa575. PubMed PMID: 32614436; PubMed Central PMCID: PMCPMC7470953.

24. Huang G, Docampo R. The mitochondrial calcium uniporter interacts with subunit c of the ATP synthase of trypanosomes and humans. mBio. 2020;11(2). Epub 2020/03/19. doi: 10.1128/mBio.00268-20. PubMed PMID: 32184243; PubMed Central PMCID: PMCPMC7078472.

25. Archer SK, Luu VD, de Queiroz RA, Brems S, Clayton C. *Trypanosoma brucei* PUF9 regulates mRNAs for proteins involved in replicative processes over the cell cycle. PLoS Pathog. 2009;5(8):e1000565. Epub 2009/08/29. doi: 10.1371/journal.ppat.1000565. PubMed PMID: 19714224; PubMed Central PMCID: PMCPMC2727004.

26. Panicucci B, Gahura O, Zikova A. Trypanosoma brucei TbIF1 inhibits the essential F1-ATPase in the infectious form of the parasite. PLoS Negl Trop Dis. 2017;11(4):e0005552. Epub 2017/04/18. doi: 10.1371/journal.pntd.0005552. PubMed PMID: 28414727; PubMed Central PMCID: PMCPMC5407850.

27. Grewal JS, McLuskey K, Das D, Myburgh E, Wilkes J, Brown E, et al. PNT1 Is a C11 cysteine peptidase essential for replication of the trypanosome kinetoplast. J Biol Chem. 2016;291(18):9492–500. Epub 2016/03/05. doi: 10.1074/jbc.M116.714972. PubMed PMID: 26940875; PubMed Central PMCID: PMCPMC4850289.

28. Klebanov-Akopyan O, Mishra A, Glousker G, Tzfati Y, Shlomai J. *Trypanosoma brucei* UMSBP2 is a single-stranded telomeric DNA binding protein essential for chromosome end protection. Nucleic Acids Res. 2018;46(15):7757–71. Epub 2018/07/15. doi: 10.1093/nar/gky597. PubMed PMID: 30007364; PubMed Central PMCID: PMCPMC6125633.

29. Aphasizheva I, Alfonzo J, Carnes J, Cestari I, Cruz-Reyes J, Goringer HU, et al. Lexis and grammar of mitochondrial RNA processing in trypanosomes. Trends Parasitol. 2020;36(4):337–55. Epub 2020/03/20. doi: 10.1016/j.pt.2020.01.006. PubMed PMID: 32191849; PubMed Central PMCID: PMCPMC7083771.

30. Schneider A. Evolution of mitochondrial protein import - lessons from trypanosomes. Biol Chem. 2020;401(6-7):663–76. Epub 2020/03/07. doi: 10.1515/hsz-2019-0444. PubMed PMID: 32142472.

31. Kaurov I, Vancova M, Schimanski B, Cadena LR, Heller J, Bily T, et al. The diverged trypanosome MICOS complex as a hub for mitochondrial cristae shaping and protein import. Curr Biol. 2018;28(21):3393–407 e5. Epub 2018/11/13. doi: 10.1016/j.cub.2018.09.008. PubMed PMID: 30415698.

32. von Kanel C, Aeschlimann S, Husova M, Oeljeklaus S, Stettler P, Schnaufer A, et al. TbTim20 facilitates protein import at a low membrane potential in trypanosomes lacking the mitochondrial genome. FEBS J. 2025. Epub 2025/10/23. doi: 10.1111/febs.70297. PubMed PMID: 41129280.

33. Niemann M, Wiese S, Mani J, Chanfon A, Jackson C, Meisinger C, et al. Mitochondrial outer membrane proteome of *Trypanosoma brucei* reveals novel factors required to maintain mitochondrial morphology. Mol Cell Proteomics. 2013;12(2):515–28. Epub 2012/12/12. doi: 10.1074/mcp.M112.023093. PubMed PMID: 23221899; PubMed Central PMCID: PMCPMC3567870.

34. Iyer A, Niemann M, Serricchio M, Dewar CE, Oeljeklaus S, Farine L, et al. The endoplasmic reticulum membrane protein complex localizes to the mitochondrial - endoplasmic reticulum interface and its subunits modulate phospholipid biosynthesis in *Trypanosoma brucei*. PLoS Pathog. 2022;18(5):e1009717. Epub 2022/05/03. doi: 10.1371/journal.ppat.1009717. PubMed PMID: 35500022; PubMed Central PMCID: PMCPMC9113592.

35. Schnaufer A. Evolution of dyskinetoplastic trypanosomes: how, and how often? Trends Parasitol. 2010;26(12):557–8. Epub 2010/08/31. doi: 10.1016/j.pt.2010.08.001. PubMed PMID: 20801716; PubMed Central PMCID: PMCPMC2932643.

36. Domingo GJ, Palazzo SS, Wang B, Pannicucci B, Salavati R, Stuart KD. Dyskinetoplastic *Trypanosoma brucei* contains functional editing complexes. Eukaryot Cell. 2003;2(3):569–77. Epub 2003/06/11. doi: 10.1128/EC.2.3.569-577.2003. PubMed PMID: 12796302; PubMed Central PMCID: PMCPMC161453.

37. Subrtova K, Panicucci B, Zikova A. ATPaseTb2, a unique membrane-bound FoF1-ATPase component, is essential in bloodstream and dyskinetoplastic trypanosomes. PLoS Pathog. 2015;11(2):e1004660. Epub 2015/02/26. doi: 10.1371/journal.ppat.1004660. PubMed PMID: 25714685; PubMed Central PMCID: PMCPMC4340940.

38. Ruhle T, Leister D. Assembly of F1F0-ATP synthases. Biochim Biophys Acta. 2015;1847(9):849–60. Epub 2015/02/11. doi: 10.1016/j.bbabio.2015.02.005. PubMed PMID: 25667968.

39. Escrivani DO, Hutchinson S, Tinti M, Wright JE, Marques CA, Faria JRC, et al. A non-coding role for trypanosome *VSG* transcripts in allelic exclusion. Nucleic Acids Res. 2025;53(19). Epub 2025/10/21. doi: 10.1093/nar/gkaf1011. PubMed PMID: 41118571; PubMed Central PMCID: PMCPMC12539627.

40. Virtanen P, Gommers R, Oliphant TE, Haberland M, Reddy T, Cournapeau D, et al. SciPy 1.0: fundamental algorithms for scientific computing in Python. Nat Methods. 2020;17(3):261–72. Epub 2020/02/06. doi: 10.1038/s41592-019-0686-2. PubMed PMID: 32015543; PubMed Central PMCID: PMCPMC7056644.

41. Ritchie ME, Phipson B, Wu D, Hu Y, Law CW, Shi W, et al. limma powers differential expression analyses for RNA-sequencing and microarray studies. Nucleic Acids Res. 2015;43(7):e47. Epub 2015/01/22. doi: 10.1093/nar/gkv007. PubMed PMID: 25605792; PubMed Central PMCID: PMCPMC4402510.

42. Tinti M. OligoSeeker. 105281/zenodo15011916. 2025.

43. Molder F, Jablonski KP, Letcher B, Hall MB, van Dyken PC, Tomkins-Tinch CH, et al. Sustainable data analysis with Snakemake. F1000Res. 2021;10:33. Epub 2021/05/29. doi: 10.12688/f1000research.29032.3. PubMed PMID: 34035898; PubMed Central PMCID: PMCPMC8114187.

44. Tinti M. myRNA-SEQ. https://githubcom/mtinti/myRna-seq. 2025.

